# Heterogeneous collateral effects in daptomycin-resistant *E. faecalis*

**DOI:** 10.1101/2023.12.07.570714

**Authors:** Anh Huynh, Jeff Maltas, Kevin B. Wood

**Affiliations:** Dept of Biophysics, University of Michigan, Ann Arbor, MI, USA; Dept of Physics, University of Michigan, Ann Arbor, MI, USA

## Abstract

Daptomycin, a cyclic lipopeptide antibiotic that targets the cell membrane, is an important therapeutic option for treating multi-drug-resistant infections, including vancomycin-resistant enterococci (VRE). Recent work has uncovered an array of daptomycin resistance mechanisms in enterococci, but relatively little is known about how these molecular defenses contribute to collateral effects–that is, to increased resistance or sensitivity to other drugs. In this work, we investigate collateral effects that arise during daptomycin adaptation of *E. faecalis* in four independent laboratory-evolved populations. Using a combination of growth assays and both single isolate and population sequencing, we identified DAP-resistant lineages with mutations in one or more genes previously associated with DAP resistance, and these isolates are characterized by divergent phenotypic properties–including different levels of DAP resistance and different growth rates (i.e. fitness costs) in drug-free media. Interestingly, we also observed strongly divergent collateral responses to different antibiotics, particularly CRO, with collateral resistance arising in mutants harboring DAP-resistance mutations in cardiolipin synthetase (*cls*) or in genes linked to the two-component signaling system YxdJK (*bceR* or a regulated transporter *ycvR*). By contrast, mutations in *liaX*, a component of a LiaFSR two-component signaling system, arose in two of the four populations, with point mutations associated with CRO-sensitivity and a large structural integration of plasmid pTEF3 associated with extreme CRO-sensitivity and a dramatically reduced growth rate. Our results reveal considerable phenotypic differences in mutations targeting the LiaSFR system and highlight trade-offs between resistance to daptomycin, collateral profiles (most notably to CRO), and drug-free growth rates in evolving lineages. As a whole, these results underscore how rich–and remarkably diverse–evolutionary dynamics can emerge even in parallel populations adapting to simple daptomycin escalation protocols.

## Introduction

Antibiotic resistance is a growing threat to global health. The development of new antimicrobial drugs is a slow and difficult process, leading to substantial recent interest in evolution-centered strategies with the potential to slow resistance based on judicious–and perhaps unorthodox–use of currently available treatments. These treatment strategies exploit interactions between cells and their environment and rely on features such as spatial heterogeneity (***Zhang et al., 2011***; ***Baym et al., 2016***; ***De Jong and Wood, 2018***; ***Feder et al., 2021***), intercellular cooperation (***Yurtsev et al., 2013***; ***Meredith et al., 2015a***,b; ***Farrokhian et al., 2022***), competition between subpopulations (***Gatenby et al., 2009***; ***West et al., 2020***; ***Hansen et al., 2020***; ***Maltas et al., 2023***), and judicious use of drug combinations simultaneously (***Zimmer et al., 2016***; ***Wood et al., 2012***; ***Torella et al., 2010***; ***Michel et al., 2008***; ***Dean et al., 2020***; ***Jahn et al., 2021***; ***Tekin et al., 2016***; ***Gjini and Wood, 2021***) or in temporal cycles (***Brown and Nathwani, 2005***; ***Bergstrom et al., 2004***; ***Nichol et al., 2015***; ***Batra et al., 2021***). One potentially promising approach is based on the concept of collateral sensitivity–an extension of so-called antagonistic pleiotropy–which, in the case of antibiotics, occurs when a population selected by one drug exhibits increased sensitivity to a different drug. Collateral effects have been a topic of substantial recent interest, with multiple studies pointing to the benefits and potential limitations of this approach (***Barbosa et al., 2019***; ***Maltas et al., 2021b***; ***Lázár et al., 2013, 2014***; ***Imamovic and Sommer, 2013***; ***Imamovic et al., 2018***; ***Roemhild et al., 2020***; ***Zhao et al., 2016***; ***Ardell and Kryazhimskiy, 2021***; ***Herencias et al., 2021***).

In this work, we offer a case-study of daptomycin resistance–and the corresponding collateral effects–in laboratory-evolved populations of *E. faecalis*, a gram-positive opportunistic pathogen that underlies a number of human infections(***Gilmore et al., 2014***). Daptomycin (DAP) is a cyclic lipopeptide antibiotic that targets the cell membrane, where it causes rapid depolarization leading to cell death (***Montero et al., 2008***; ***Pogliano et al., 2012***; ***Tran et al., 2015***; ***Taylor and Palmer, 2016***). Daptomycin is an important therapeutic option for treating multi-drug-resistant infections (***Streit et al., 2004***; ***Steenbergen et al., 2005***; ***Fowler Jr et al., 2006***; ***Yim et al., 2017***; ***de Las Revillas et al., 2018***), including vancomycin-resistant enterococci (VRE). However, daptomycin resistance in enterococcus has been increasingly reported in the last decade in both the laboratory and the clinic (***Kelesidis et al., 2011***; ***Kamboj et al., 2011***; ***Kelesidis et al., 2012***; ***Humphries et al., 2013***; ***Miller et al., 2016***; ***Kinnear et al., 2019, 2020***).

Daptomycin resistance in enterococcus has been linked with dozens of genes (***Arias et al., 2011***; ***Palmer et al., 2011***; ***Humphries et al., 2013***; ***Diaz et al., 2014***; ***Nair et al., 2023***), many associated with multi-component signaling systems such as LiaFSR, which modulates the response of the cell membrane to stress (***Tran et al., 2013, 2015***; ***Rincon et al., 2019***; ***Khan et al., 2019a***; ***Ota et al., 2021***; ***Reyes et al., 2015***; ***Ota et al., 2021***). In lab evolution experiments, mutations in the liaFSR system–or putative downstream targets–dominate the early stages of DAP-adaptation, followed by major increases in DAP resistance linked to other genes–for example, those encoding cardiolipin synthase (***Miller et al., 2013***; ***Palmer et al., 2011***). In strains lacking *liaR*, increasing doses of sub-MIC daptomycin selected for high-cost mutations in an alternative two-component signaling system (YxdJK) (***Miller et al., 2019***), which itself has been linked to bacitracin resistance ***Gebhard et al***. (***2014***) (which targets cell wall synthesis), as well as mutations in a putative dihydroxyacetone kinase (DAK) family enzyme.

Collateral effects arising from daptomycin selection–and selection by a wide range of other antibiotics and environmental stressors (***Maltas and Wood, 2019***; ***Maltas et al., 2020, 2021a***; ***Yong et al., 2023***)–have also been characterized phenotypically and, to a limited degree, genetically. At a global level–when collateral effects from all selecting drugs are considered–collateral profiles appear dynamic, with collateral resistance arising more frequently in early stages of adaptation and collateral sensitivity arising more frequently after longer periods of selection (***Maltas et al., 2021a***). At the level of individual selecting drugs, however, the patterns are idiosyncratic (***Maltas et al., 2021a***), with some collateral effects matching the global trend (e.g. populations selected by linezolid show increased collateral sensitivity to ceftriaxone (CRO) over time) while other populations show opposing trends (e.g. populations selected by ciprofloxacin show decreasing collateral sensitivity to CRO over time).

In this work, we investigate how collateral effects arise in *E. faecalis* populations selected by daptomycin in laboratory evolution experiments spanning approximately 60 generations. Previous work has shown that isolates selected in daptomycin are characterized by heterogeneous collateral profiles (***Maltas and Wood, 2019***; ***Maltas et al., 2021a***). Most notably, the collateral response to ceftriaxone (CRO), an inhibitor of cell wall synthesis, ranges from resistant to highly sensitive in different isolates. The latter is reminiscent of the so-called “seesaw” effect, an inverse relationship between MICs to glycopeptides or lipopeptides and beta-lactams commonly observed in other Gram-positive species (***Sieradzki and Tomasz, 1999***; ***Sieradzki et al., 2003***; ***Yang et al., 2010***; ***Renzoni et al., 2017***; ***Barber et al., 2014***). A similar “seesaw” effect has also been observed in *E. faecalis* (***Sakoulas et al., 2013***; ***Jahanbakhsh et al., 2020***; ***Maltas and Wood, 2019***), and recent work has linked CRO hypersensitivity to DAP-driven mutations in the LiaSFR system (***Khan, 2020***; ***Khan et al., 2019b***). To examine how these collateral effects arise during daptomycin adaptation, we phenotypically and genetically characterized dozens of individual isolates selected from each of four daptomycin-selected populations (***Maltas et al., 2021a***). In all populations, we identified isolates with mutations in one or more genes previously associated with DAP resistance, and these isolates are characterized by divergent phenotypic properties–including different levels of DAP resistance and different growth rates (i.e. fitness costs) in drug-free media. Interestingly, we also observed strongly divergent collateral responses to different antibiotics, particularly CRO, with collateral resistance arising in mutants harboring DAP-resistance mutations in cardiolipin synthetase (*cls*) or in genes linked to the two-componenent signaligng sytem YxdJK (*bceR* or *ycvR*) (***Miller et al., 2019***). By contrast, mutations in *liaX* arose in two of the four populations, with point mutations associated with CRO-sensitivity and a large structural integration of plasmid pTEF3 associated with extreme CRO-sensitivity and a dramatically reduced growth rate. Our results reveal that multiple daptomycin-resistant lineages can emerge even in simple lab evolution protocols, reveal considerable phenotypic differences in mutations targeting the LiaSFR system, and highlight trade-offs between resistance to daptomycin, collateral profiles (most notably to CRO), and drug-free growth rates (fitness costs) in evolving lineages.

## Results

### Heterogeneous daptomycin resistance in DAP-selected populations

To investigate daptomycin resistance in the lab, we focused on four daptomycin-adapted populations described in (***Maltas et al., 2021a***). Briefly, we exposed four independent populations (each from a single colony of the ancestral V583 strain) to increasing concentrations of daptomycin over 8 days with 1:200 daily dilutions (Figure 1). Each day, 3 vials–each at a different DAP concentration, with concentrations spanning sub- and super-MIC levels–were seeded with dilutions from the previous day’s population. At the end of the day, the sample that survived in the highest concentration was propagated on to the next day, where 3 new concentrations were chosen (typically one-half, 2x and 4x the highest concentration with growth from the previous day; see Methods). Despite using a similar protocol to increase DAP concentration in each adaptation experiment, the variable responses of the 4 populations led to distinct differences in the DAP survival concentration (defined here as the highest concentration of DAP for which a given lineage survived each day) over time (Figure 1). In previous work, single isolates from these populations, along with those adapted to other drugs, were used to characterize collateral effects to a large library of antibiotics and other stressors (***Maltas and Wood, 2019***; ***Maltas et al., 2020, 2021a***). Among a collection of collateral profiles, the single isolates selected from these four DAP-selected populations stood out as being particularly heterogeneous and potentially dynamic over time.

**Figure 1.**
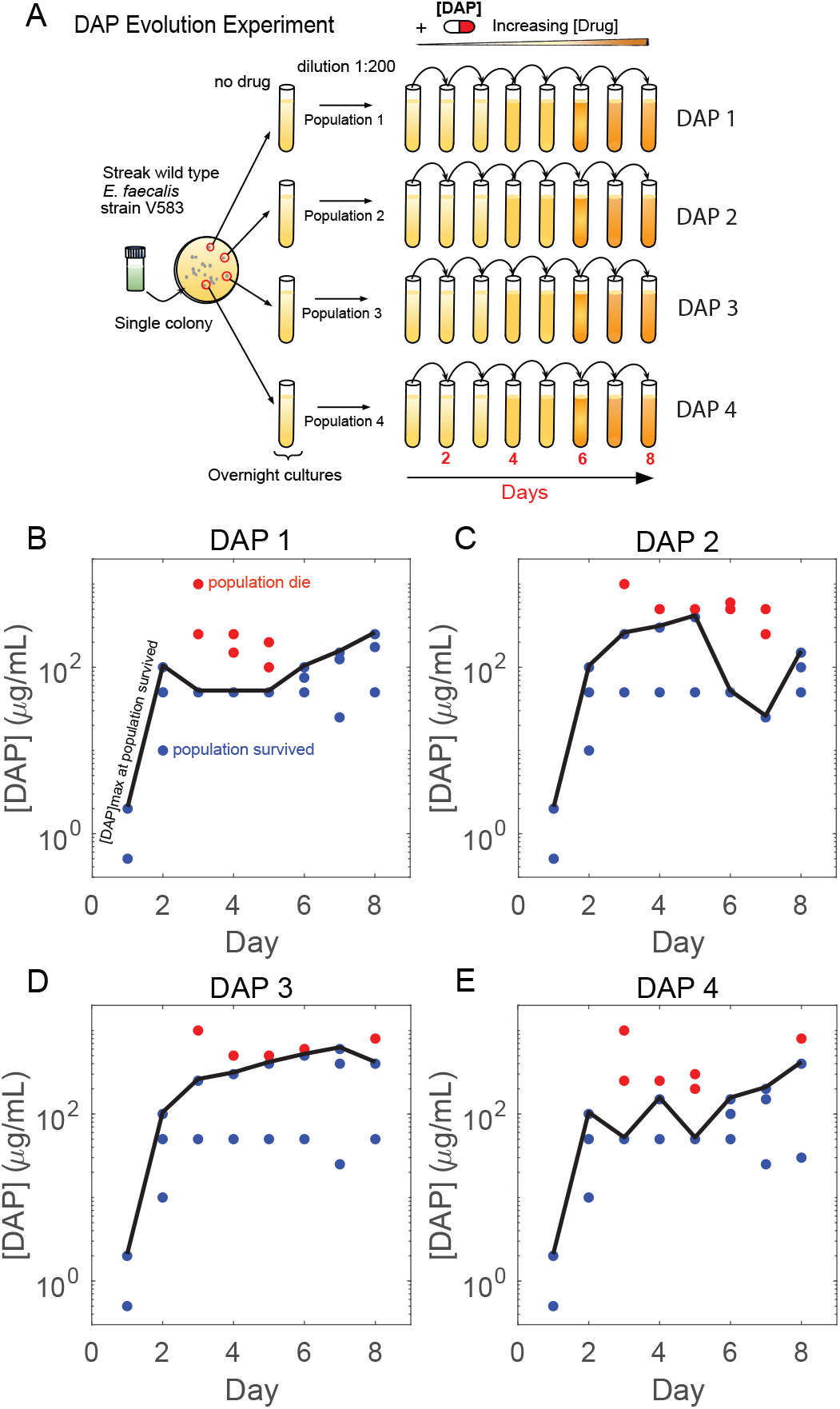
Evolution of daptomycin resistance in the lab. A) Schematic of laboratory evolution experiment with *E. faecalis* strain V583 treated with Daptomycin (DAP). The overnight cultures were exposed to (typically) increasing concentrations over an 8-day serial-passage with daily dilutions of 1:200. B) Panels B to E show trajectories of the 4 DAP populations; black curves represent the survival concentration (defined here as the highest concentration of DAP for which a given lineage survived each day); blue (red) circles represent viable (non-viable) populations, as inferred from turbidity (see Methods, Chapter 2).

To characterize these adapting lineages, we isolated 12 single colonies from each of the four DAP populations at 4 different time points (days 2, 4, 6, and 8) and estimated the IC_50_ to DAP for each isolate from dose response curves (measured in technical replicates of 4). Despite similar evolution protocols, we found that resistance to DAP varied across the four populations (Figure 2). In three of the populations (DAP1, DAP2, DAP4), we observed substantial DAP resistance by day 2, while isolates from DAP3 did not achieve comparable levels of resistance until day 6. In addition, the distribution of IC_50_’s varies across time, within individual populations, as well as across populations. Interestingly, these distributions appear bimodal at several time points, suggesting that multiple resistant lineages may be present–at substantial fractions– simultaneously.

**Figure 2.**
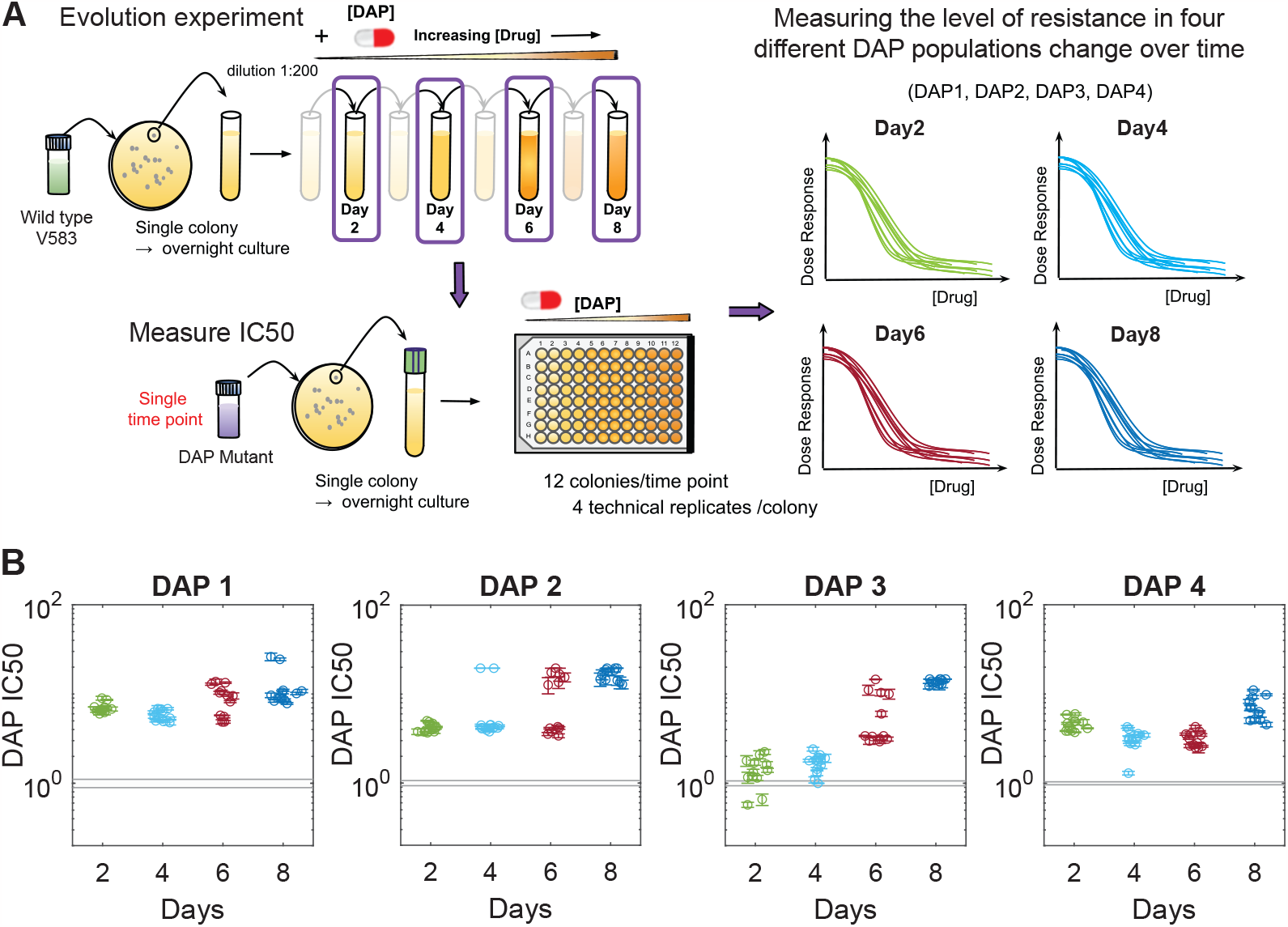
Daptomycin resistance is heterogeneous and dynamic within individual populations. A) Dose-response curves (in technical replicates of 4) for twelve isolates from each DAP-evolved population (DAP-1 to DAP-4) at four different time points (days 2, 4, 6, and 8) were used to estimate IC_50_ to daptomycin. B) Daptomycin IC_50_ for each isolate, normalized by the IC_50_ of a collection of ancestral populations measured on the same day, over time for each of the four populations. Shaded region shows precision (± 3 standard error) of IC_50_ estimates in the ancestral control isolates. Isolates above the shaded region are considered DAP-resistant, while those below the shaded region exhibit increased DAP-sensitivity. Individual points represent the mean IC_50_ (± standard error from 4 technical replicates) for a single colony.

### Collateral effects to CRO vary dramatically across and within populations

Given the range of DAP-resistance in isolates, we next asked how the collateral effects to CRO change within and across populations. To estimate the distribution of collateral effects, we measured dose response curves to CRO for each of the previously selected isolates along with 36 additional isolates from each of the four time points for population DAP-1. Indeed, we found substantial variability in CRO sensitivity across isolates (Figure 3). In two of the four populations (DAP3 and DAP4), we observed low-levels of CRO resistance by day 2, with the distribution shifting slightly towards increased CRO resistance over time. By contrast, a substantial fraction of isolates in populations DAP1 and DAP2 were collaterally sensitive to CRO on day 2. In DAP-1, the distribution drifts to higher CRO resistance levels over time, though a subpopulation of highly CRO sensitive isolates– with decreases in IC_50_ of nearly 100 fold relative to the ancestral strain–emerges and is particularly noticeable by day 8. Surprisingly, the DAP2 population is dominated by the hyper-sensitive isolates even on day 2, and collateral sensitivity remains high throughout. These results suggest that the evolution of collateral effects in response to DAP exposure exhibits rich dynamics, even on relatively short timescales.

**Figure 3.**
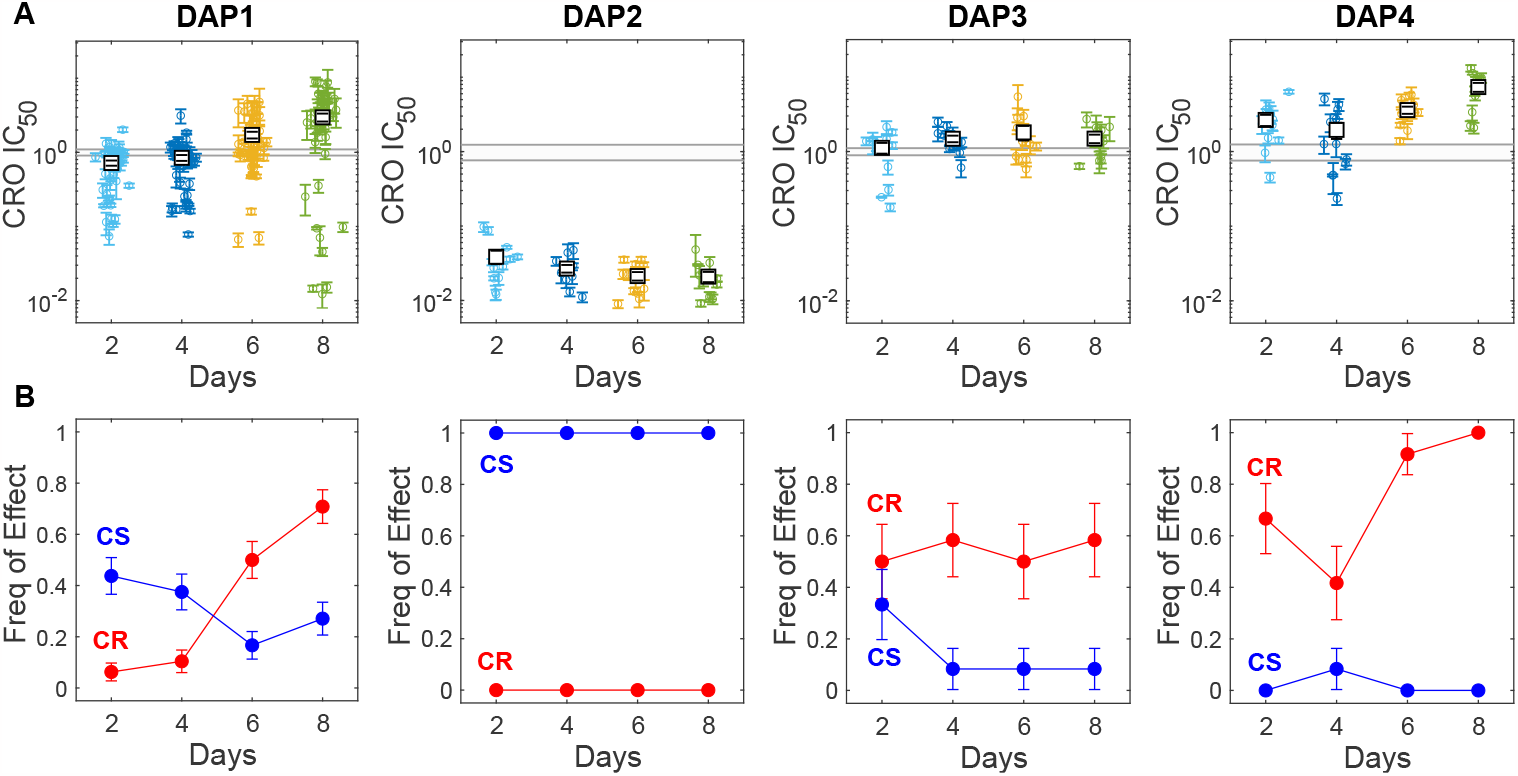
Collateral CRO profiles differ markedly across and within DAP-selected populations. Dose-response curves (in technical replicates of 4) for 48 isolates at four different time points (days 2, 4, 6, and 8) from population DAP1 along with 12 isolates at four time points for populations DAP2-DAP4 were used to estimate IC_50_ to ceftriaxone (CRO). A) CRO IC_50_ for each isolate, normalized by the CRO IC_50_ of a collection of ancestral populations measured on the same day, over time for each of the four populations. Shaded region shows precision (± 3 standard error) of IC_50_ estimates in the ancestral control isolates. Isolates above the shaded region are considered CRO-resistant, while those below the shaded region exhibit increased CRO-sensitivity. Individual points represent the mean IC_50_ (± standard error from 4 technical replicates) for a single colony. B) Frequency of collateral effects in individual isolates over time in four different DAP populations, with collateral sensitivity (CS) represented in blue and collateral resistance in red (CR).

### Idiosyncratic covariation of DAP and CRO resistance across populations

We also measured the covariation of DAP and CRO resistance in a subset of individual isolates from the four DAP-selected populations (Figure 4). We found a range of different dynamics in the 2-d space of IC_50_s. Populations DAP1 and DAP4 are characterized by IC_50_ distributions whose mean drifts towards increased resistance to both drugs over time, though DAP1 includes a highly CRO-sensitive subpopulation by day 8. DAP2 isolates move toward increasing DAP resistance and decreasing CRO resistance, while DAP3 isolates show increasing DAP resistance but relatively little change in CRO resistance (which is low-level collateral resistance) over time.

**Figure 4.**
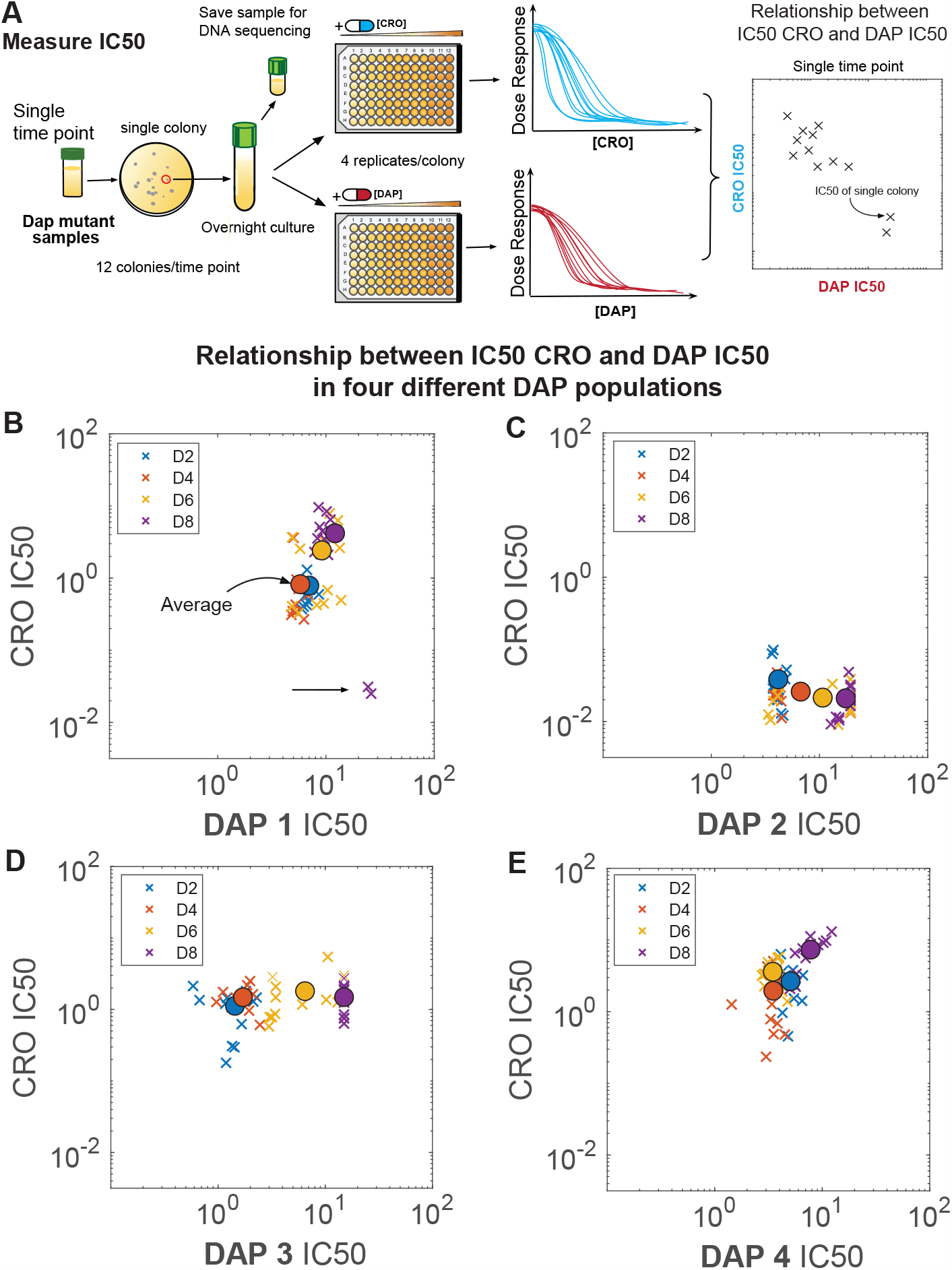
Co-variation of DAP and CRO resistance across populations. A) Schematic of measuring covariation in CRO and DAP resistance in four different DAP-adapted populations. For each time point, dose-response curves for both CRO and DAP (each in technical replicates of 4) were measured for 12 isolates. B) IC_50_s for each drug are normalized by IC_50_ values in a collection of ancestral isolates measured on the same day. The IC_50_ of a single colony is represented by an x, while the population average of 12 colonies is represented by a circle. Isolates from day 2 are in blue, from day 4 are in orange, from day 6 are in yellow, and from day 8 are in purple.

### Population and single-isolate genome sequencing reveals varied combinations of mutations in genes associated with DAP resistance

To investigate the genetic underpinnings of the varied daptomycin-resistance dynamics, we performed population sequencing on each of the four populations over time and also sequenced the full genomes of 40 individual isolates. Despite the simplicity of the laboratory evolution, we observed a surprising diversity of DAP-R mutations across different populations (Figure 5). We identified a number of mutations in genes previously linked to DAP resistance (Table 1), and we found that different combinations of these mutations accumulated over time in different populations.

**Table 1.**
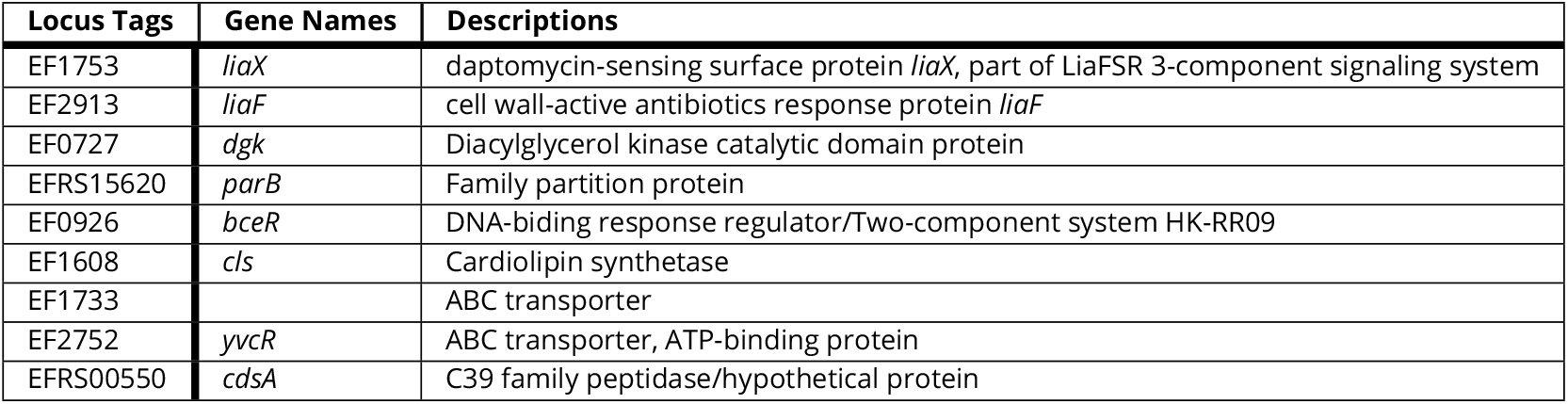
Mutations identified in selected populations.

**Table 2.**
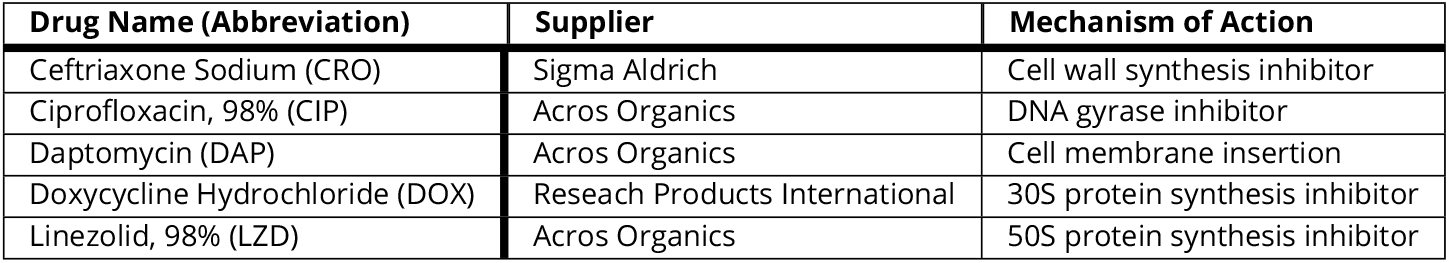
Antibiotics used in this study.

| Drug Name (Abbreviation) | Supplier | Mechanism of Action |
| --- | --- | --- |
| Ceftriaxone Sodium (CRO) | Sigma Aldrich | Cell wall synthesis inhibitor |
| Ciprofloxacin, 98% (CIP) | Acros Organics | DNA gyrase inhibitor |
| Daptomycin (DAP) | Acros Organics | Cell membrane insertion |
| Doxycycline Hydrochloride (DOX) | Research Products International | 30S protein synthesis inhibitor |
| Linezolid, 98% (LZD) | Acros Organics | 50S protein synthesis inhibitor |

**Table 3.**
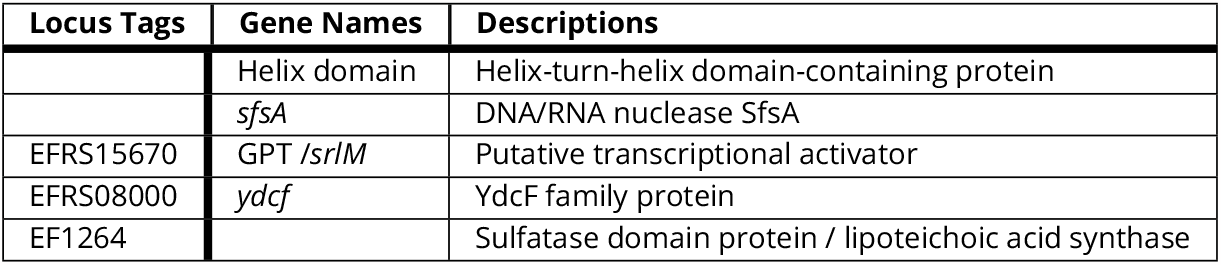
Additional mutations observed in population sequencing.

**Figure 5.**
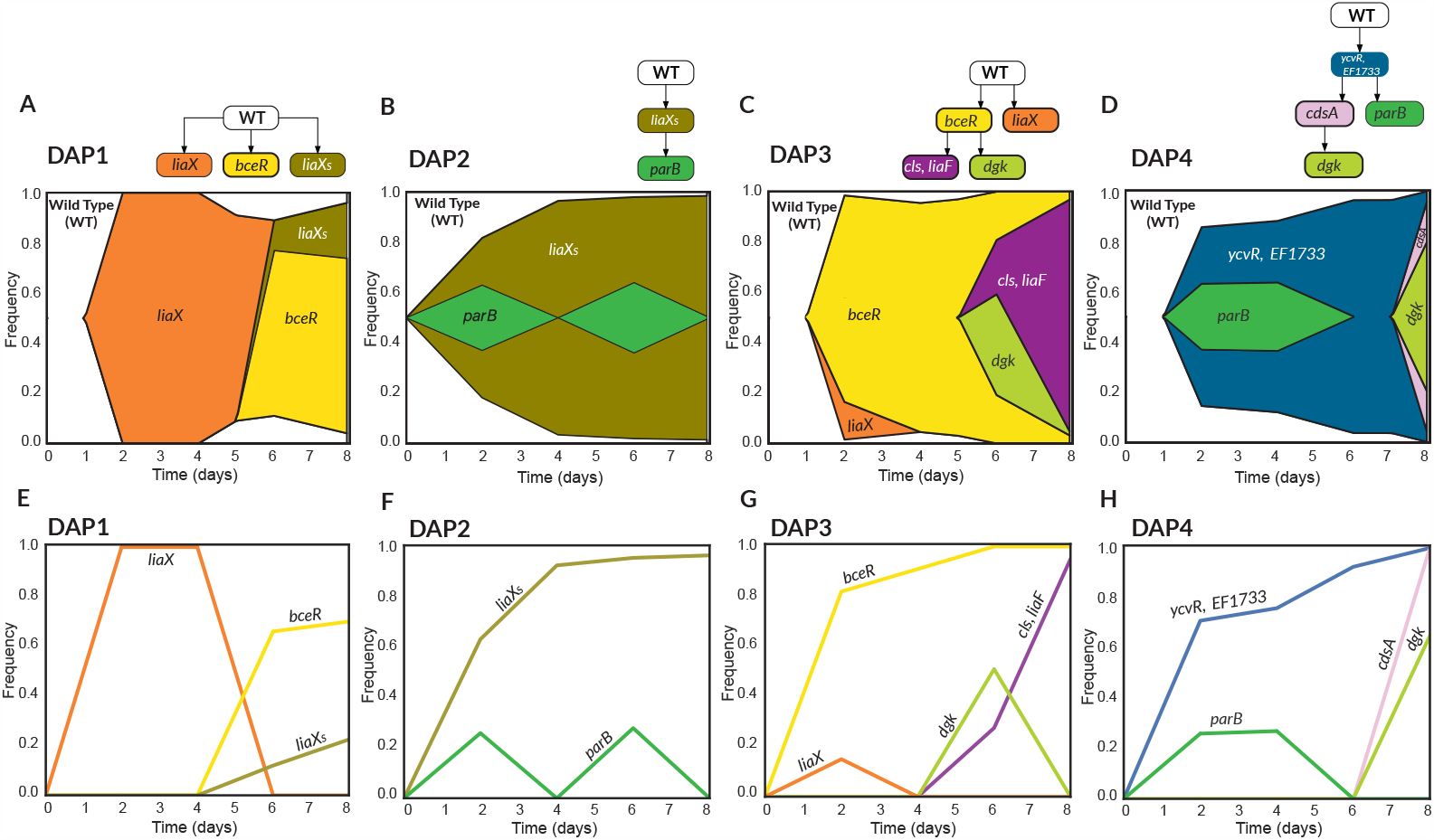
Evolutionary dynamics of V583 evolving Daptomycin through independent evolution experiments. Panels A–D correspond to the genotype tree and Muller plots for each DAP-adapted population over time. Panels E-H display individual genotype frequencies corresponding to each Muller plot. Muller plots were calculated from population sequencing data using the Lolipop pipeline, which is described in (***Scribner et al., 2020***) and available on GitHub. See also Table 1 for a description of different mutations and Figure S1.

In three of the four populations (DAP1, DAP2, DAP3), mutations in the liaFSR occurred within the early stages of adaptation, consistent with previous findings (***Miller et al., 2013***; ***Palmer et al., 2011***). Point mutations in the gene *liaX* swept (nearly) to fixation in population 1 but occurred only at low frequency in DAP3. We also identified larger structural mutations in *liaX* in DAP1 and DAP2– and in the latter case, this structural mutant (which we refer to as liaXs) swept to fixation over time, while it remained a low-frequency variant in DAP1.

We also observed multiple DAP-R mutations not directly linked with liaFSR. In populations DAP1 and DAP3, lineages characterized by point mutations in *bceR*, a gene associated with the two-component signaling system YxdJK previously linked with resistance to bacitracin and found in DAP-adapted strains lacking the liaFSR system (***Miller et al., 2019***), appeared to out-compete the initial liaX mutants. In DAP3, these *bceR* lineages split into two competing lineages, one that acquired an additional mutation in *dgk*, a gene encoding a putative kinase involved in lipid metabolism, and a second with mutations in both *liaF* (from the liaFSR system) and *cls*. To our knowledge, the mutation in *dgk* has not been reported elsewhere, but it is reminiscent of similar adaptations previously seen in a dihydroxyacetone kinase (DAK) associated with lipid metabolism (***Miller et al., 2019***). Mutations in cardiolypin synthetases, such as those in the *cls* gene, have also been commonly linked with DAP resistance (***Palmer et al., 2011***). Finally, in DAP4, a lineage dominated by a double mutant– with mutations in *ycvR* and EF1733 (a putative ABC transporter)– rose to high frequencies by day 6. This lineage later acquired mutations in *cdsA*, a C39 family peptidase linked with DAP resistance in other Streptococcus species (***Tran et al., 2019***), and subsequently in *dgk*. Finally, we observed mutations in *parB*, associated with chromosome or plasmid segregation but not, to our knowledge, with DAP resistance, at 4 different time points (days 2 and 6 of DAP2; days 2 and 4 of DAP4). While the pattern of their appearance, as well as the fact that these variants did not appear in any of the 40 individual isolates, suggests they may be spurious variant calls, we keep them here for completeness and because they appeared in multiple populations.

### DAP-R mutants exhibit diverse range of resistance levels and fitness costs

The genome sequencing reveals a surprisingly complex collection of evolutionary trajectories that include different combinations of individual mutations. However, it is not clear how a given combination of mutations are related to phenotypic properties of a given mutant. To investigate this question, we measured drug-free growth rate as well as DAP- and CRO-resistance (IC_50_) levels for isolates representing seven different mutant “classes” which were found among the 40 single-colony isolates. These mutants included single gene mutants (mutations in *bceR* and *liaX*), 2 double mutants (mutations in *dgk* and *bceR*; or in EF1733 and *ycvR*), and 2 triple mutants (mutations in *bceR, cls*, and *liaF*; or in *bceR, cls*, and *liaF*).

We found that the different mutant classes exhibit considerable differences in drug-free growth rate (referred to here at “growth cost”, see Methods), direct resistance levels to DAP, and collateral resistance levels to CRO (Figure 6; see also Figures S3 and S4). DAP resistance is weakly correlated with growth cost (Figure 6, left panel), with a double (*dgk, bceR*) and a triple (*bceR, cls, liaF*) mutant from DAP3 showing the highest levels of DAP resistance and elevated costs, while single mutants (*bceR* and *liaX*) show lower levels of DAP-R but also decreased cost. In addition, all mutations outside of *liaX* showed small to moderate levels of collateral resistance to CRO. By contrast, mutations in *liaX* were associated with collateral sensitivity.

**Figure 6.**
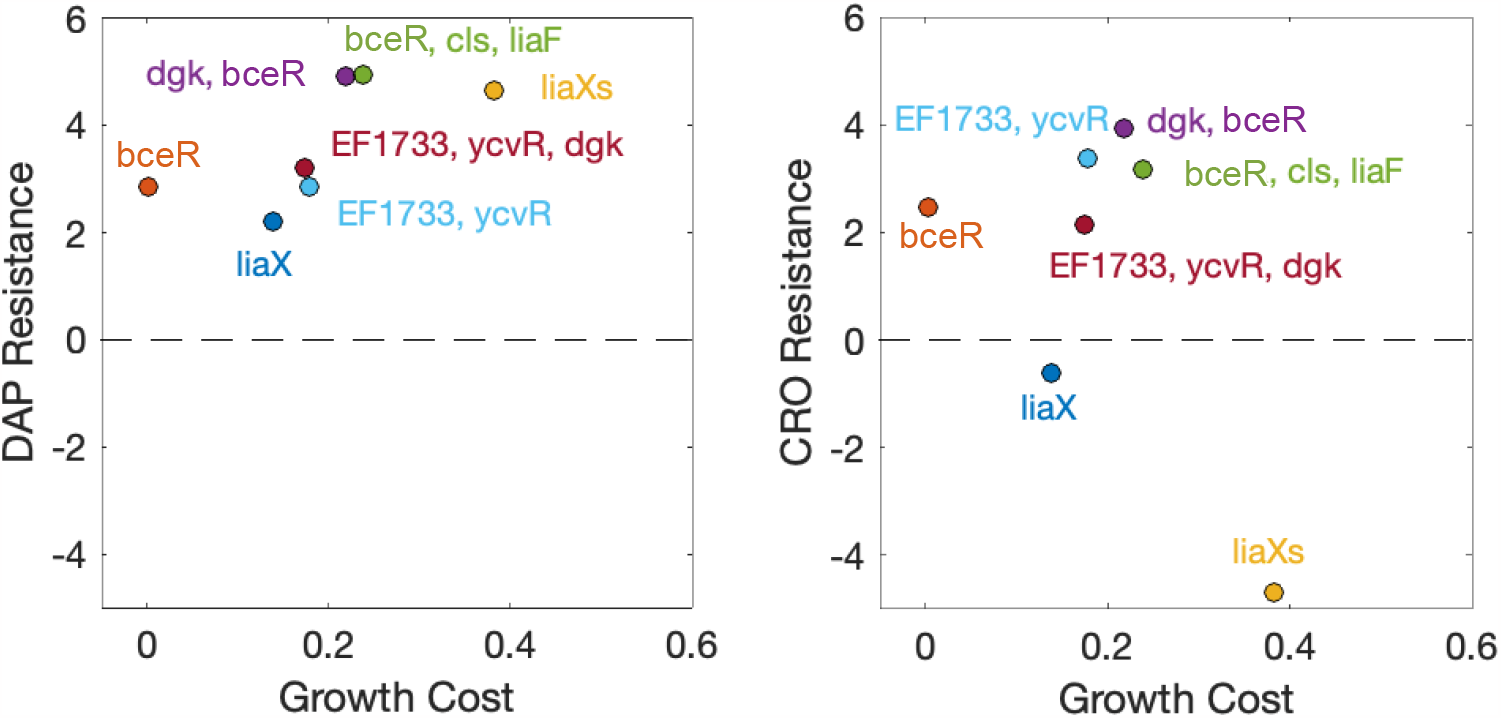
Mutations and growth costs in isolates from different DAP-adapted populations. DAP resistance (left) and CRO resistance (right) vs growth cost for isolates representing each of the 7 observed mutants. Resistance is defined as the (log2-transformed) fold change in IC_50_ relative to the ancestral strain; positive values represent increased resistance, while negative values represent increased sensitivity. Growth cost is defined as the fractional decrease in maximum growth rate when strains are grown in drug-free media. The mutants include 3 single gene mutants (mutations in *bceR* and two different mutations in *liaX*, a single nucleotide variant, and a pTEF3 insertion variant, the latter of which we designate by using an “s” suffix), 2 double mutants (mutations in *dgk* and *bceR*; or in EF1733 and *ycvR*), and 2 triple mutants (mutations in *bceR, cls*, and *liaF*; or in *bceR, cls*, and *liaF*).

### V583 plasmid pTEF3 integrates into *liaX* and confers CRO hypersensitivity

The short read sequencing identified likely structural changes in *liaX* in the DAP2 mutant. However, short read sequencing is often inadequate for the assessment of structural changes. Therefore we performed additional ONT long reads sequencing which allowed for near-perfect genome assemblies when used in combination with the short reads (Methods and Materials) (***Wick et al., 2023***). Briefly, we assembled the genomes using exclusively ONT long reads and followed this with long-read polishing. This allows for an accurate large-scale genome assembly, likely containing only small scale errors (such as single-bp substitutions, deletions, insertions). We then used the Illumina sequencing short read to polish these small scale errors away.

Remarkably, the full genome assembly revealed an integration of V583’s complete pTEF3 plasmid, flanked by two copies of a IS1216-like transposase, into *liaX* at amino acid position 241 (Figure S5). These findings are complimentary to recent work that found a frameshift stop codon at amino acid position 289 which resulted in truncation of the C-terminal domain and conferred a daptomycin-resistant phenotype (***Khan et al., 2019a***) and, separately, to recent work indicating that phage or antibiotic stress can drive rapid genome-scale transposition in enterococci, including in *E. faecium* from human patients treated with daptomycin (***Kirsch et al., 2023***).

In comparison to the single nucleotide variant, pTEF3 integration dramatically increased the resistance to DAP as well as the growth cost in drug-free media (Figure 6, left panel). Interestingly, pTEF3 integration into *liaX* conferred CRO hypersensitivity (Figure 6, right panel), a finding that is consistent with work linking CRO hypersensitivity to DAP-driven mutations in the LiaSFR system (***Khan, 2020***; ***Khan et al., 2019b***).

### Disparate growth costs and resistance levels favor different mutants at low and high DAP concentrations

In population DAP1 at days 6 and 8 we observed coexisting mutants (mutation in *bceR* and pTEF3 insertion in *liaX*, which we denote with subscript “s”) with substantial differences in DAP IC_50_s and growth costs (Figure 5 and Figure 6). To further characterize these competing lineages, we measured growth rates of each at multiple concentrations of DAP as well as 3 additional antibiotics (DOX, CIP, LZD). Consistent with the previous end-point IC_50_ measurements (Figure 6), we found that the two mutants exhibit considerable differences in growth rate as function of DAP, with the bceR mutant showing increased growth at low DAP concentrations–a result, in part, of its smaller fitness cost–while the liaXs mutant has a growth advantage at DAP concentrations greater than about 40 ug/mL (Figure 7).

**Figure 7.**
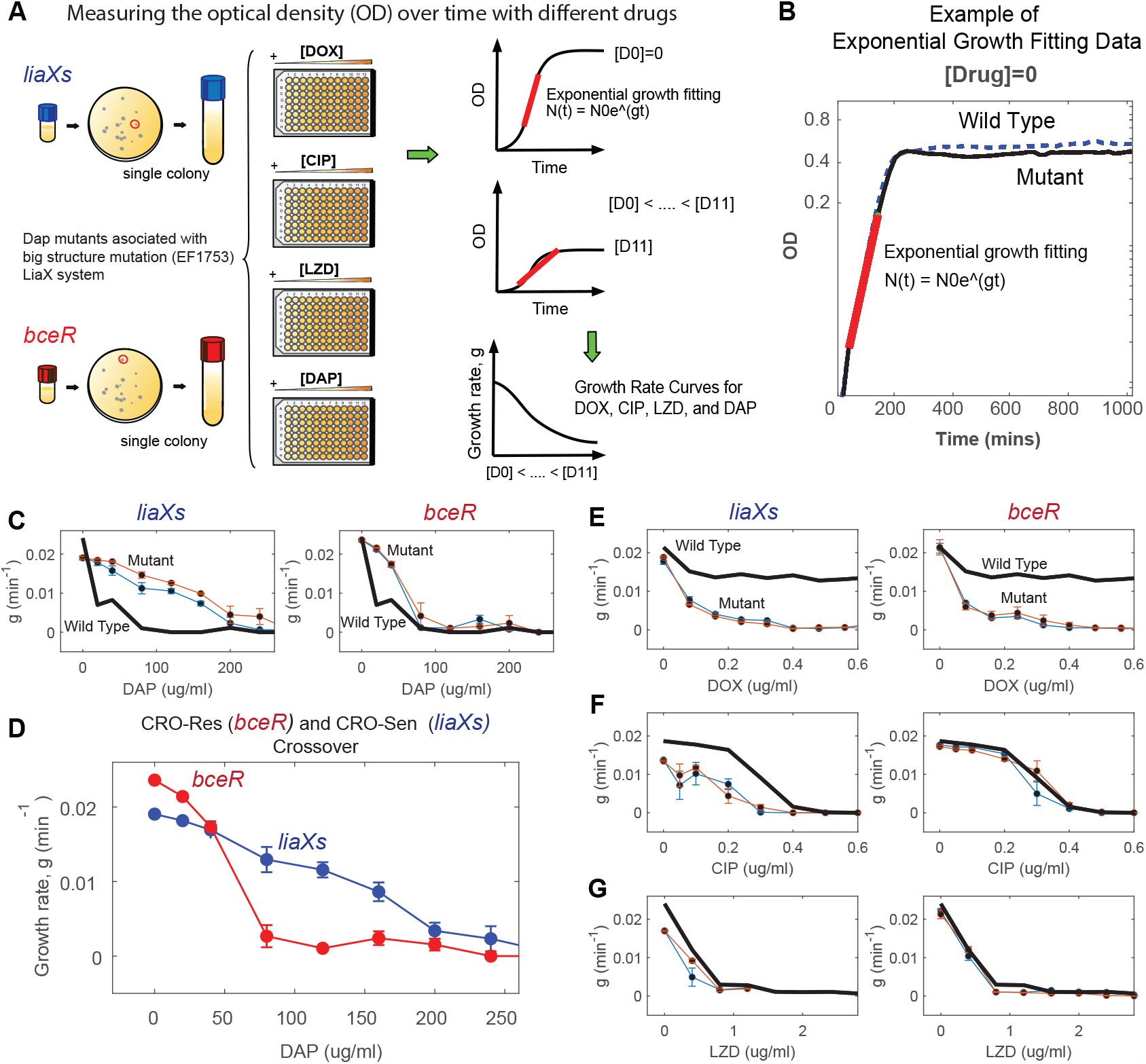
Effects of different antibiotics on growth rates of *liaX*_*S*_ and *bceR* mutants. A) Schematic of the measurement of per capita growth rate from optical density time series in two isolates isolated from DAP1 populations. The first isolate has a pTEF3 plasmid insertion in EF1753 (denoted *liaX*_*S*_); the second isolate has a point mutation in a repressor (*bceR*) linked to the regulatory system YxdJK. B) Per capita growth rate was estimated from time series of optical density in exponential phase using nonlinear least squares fitting. Red line: example growth rate fits for two isolates with similar growth rates (blue and black curves). C) Per capita growth rate as a function of daptomycin (DAP). Black curves show growth of ancestral (WT) population. Individual points are the means over 4 technical replicates, with errorbars plus/minus 1 standard error. D) Growth rate dose-response curves are characterized by low-DAP regimes where *bceR*-mutant grows faster and high-DAP regimes where *liaX*_*S*_ -mutant grows faster. The crossover point occurs at a DAP concentration of approximately 40 ug/ml. E-G) Same as C, per capita growth rate as a function of doxycycline (DOX, upper panel), ciprofloxacin (CIP, middle panels), and linezolid (LZD, lower panels) for *liaX*_*S*_ -mutant (left column) and *bceR*-mutant (right column).

On the other hand, the bceR mutant tends to show increased growth (relative to the liaXs mutant) in the presence of other drugs, consistent with the opposing collateral effects we saw for CRO based on end-point measurements. While growth of bceR mutants in the presence of CIP and LZD is similar to that of WT cells over most concentrations tested, the increased growth cost associated with liaXs leads to a substantial reduction in growth at low levels of CIP and LZD (Figure 7). In addition, both mutants are strongly sensitized to doxycycline, with growth rates nearing zero at concentrations largely sub-inhibitory in the wild type (ancestral) cells.

## Discussion

Daptomycin is one of the last defenses against infections caused by multi-drug resistant bacteria, and the increasing prevalence of daptomycin resistance poses obstacles to the effective treatment of these infections. The results reported here add to our growing understanding of the molecular and genetic underpinnings of daptomycin resistance. Despite the relatively simple adaptation protocol–one performed under well-controlled laboratory conditions that are substantially simplifications of the natural environments faced in vivo–the evolution of daptomycin resistance is surprisingly complex and, in four independent populations, followed four largely distinct evolutionary pathways to resistance.

Our results complement, and are consistent with, a number of studies on daptomycin resistance that identified most of the mutational targets we observed ***Arias et al***. (***2011***); ***Palmer et al***. (***2011***); ***Humphries et al***. (***2013***); ***Diaz et al***. (***2014***); ***Tran et al***. (***2013***, 2015); ***Rincon et al***. (***2019***); ***Khan et al***. (***2019a***); ***Ota et al***. (***2021***); ***Reyes et al***. (***2015***); ***Miller et al***. (***2013***); ***Nguyen et al***. (***2023***); ***Miller et al***. (***2023***); **?**. Collateral effects have been explored in much less detail, though the collateral sensitivity we observe is consistent with the “seesaw” effect observed with glycopeptides or lipopeptides and beta-lactams in other Gram-positive species ***Sieradzki and Tomasz*** (***1999***); ***Sieradzki et al***. (***2003***); ***Yang et al***. (***2010***); ***Renzoni et al***. (***2017***); ***Barber et al***. (***2014***). A similar “seesaw” effect was recently reported in *E. faecalis* ***Sakoulas et al***. (***2013***); ***Jahanbakhsh et al***. (***2020***); ***Maltas and Wood*** (***2019***). Most notably, recent work is offering an increasingly clear picture of membrane adaptation in daptomycin-resistant enterococci ***Nguyen et al***. (***2023***) and has linked CRO hypersensitivity to DAP-driven mutations in the LiaSFR system ***Khan*** (***2020***); ***Khan et al***. (***2019b***). We are hopeful that their continued efforts will lead to a deeper understanding of the molecular basis of this collateral sensitivity.

Collateral effects have also been reported in isolates of a related enterococcal species (*E. faecium*), which acquired sensitivity to other drugs when evolved in the lab in the presence of daptomycin ***Zeng et al***. (***2022***). They found daptomycin resistant mutations associated with the liaFSR system and *yycFG* and *cls* genes, all of which were associated with fitness costs. Additionally, they observed daptomycin resistant mutants with increased collateral sensitivity to the glycopeptide antibiotic vancomycin due to a large-scale insertion mediated by the IS1216E-transposon. Our results include a similar structural insertion–this time the pTEF3 plasmid in *liaX*–and similar insertion events have been recently observed in *E. feacalis* populations under phage stress and in *E. faecium* from daptomycin-treated patients ***Kirsch et al***. (***2023***), suggesting that transposon-mediated dynamics play an important role in maintaining genome plasticity of enterococci. Recently developed sequence-based technologies offer promising approaches to interrogate these insertion sequence dynamics in more detail (***Kirsch et al., 2023***). In addition, the insertion sequence IS1216 been previously observed flanking resistance genes in *E. faecalis* ***Li et al***. (***2023***).

Similar results–most notably, mutations involving the cls gene and those associated with the LiaSRF system ***Zeng et al***. (***2022***); ***Arias et al***. (***2011***); ***Woods et al***. (***2023***)–have also been observed in the clinic. In one example, researchers isolated strains of *E. faecalis* from a patient with fatal bacteremia before and after treatment with DAP. They found mutations associated with the liaF gene, cls, and GdpD (an enzyme also involved in phospholipid metabolism) ***Arias et al***. (***2011***). Similarly, Woods and colleagues ***Woods et al***. (***2023***) found daptomycin resistance in *E. faecium* can evolve via multiple separate evolutionary pathways in patients, with most strains harboring mutations associated with the cardiolipin synthase (clsA) gene. Daptomycin resistance was also shown to arise in off-target gut-associated populations during intravenous treatment ***Kinnear et al***. (***2020***).

Our results illustrate how DAP-R mutations can arise in different combinations, even under similar environmental pressures, and lead to different effects of fitness and collateral profiles. The work highlights several challenges, and potential opportunities, for applying collateral sensitivity in the clinic. In particular, it highlights that steering evolution down particular daptomycin-resistant pathways–in this case, pathways targeting the *liaX* gene–may lead to strong collateral sensitivities, opening the door to drug-switching strategies that target populations at their most vulnerable. On the other hand, we also showed that treatment with daptomycin under very similar conditions led to highly variable evolutionary trajectories across replicate populations. It is not clear the extent to which this variability is specific to daptomycin, which targets that cell membrane and may therefore be susceptible to a wide range of resistance mechanisms that module membrane composition and homeostasis. In addition, our adaptation experiments were performed in relatively small populations–potentially much smaller than those in the gut of human hosts. Previous work in chemostats ***Miller et al***. (***2013***, 2019)–using larger populations–has indeed shown that daptomycin resistance occurs via a relatively predictable sequence, and it would be interesting to probe daptomycin adaptation at intermediate population sizes between these two extremes. In any event, harnessing these (apparently) stochastic trajectories may require application of advanced controltheoretic approaches, some of which have been recently applied to steering evolution ***Maltas and Wood*** (***2019***); ***Iram et al***. (***2020***). Future work may be able to exploit the fitness costs associated with certain genes to design more effective control strategies to steer evolution along clinically beneficial trajectories.

## Methods and Materials

### Strains, growth conditions, and drugs

Evolution experiments were conducted using the *E. faecalis* strain V583 and the resistance was derived from the same strain. Samples were stocked in 30% glycerol and stored at -80 C. Cultures for experiments were taken from single colonies grown on agar plates and then inoculated at 37^°^ C overnight before dilution in fresh media and the beginning of the experiment. All experiments and dose-response measurements were conducted in Brain-Heart Infusion or BHI (Remel). Drug stock solutions were prepared from powdered stock and stored at −20^°^ C in single-use aliquots. All experiments with daptomycin were supplemented with 20 *μ*g/mL Ca^2+^.

### Measuring drug resistance

To measure IC_50_, a sample acquired from the evolution experiment was streaked on a BHI Agar plate. From there, a single colony was selected and used for inoculation and incubated at 37° C overnight. Cells were then exposed to a drug concentration gradient, differing among each of the 12 wells on the plate, and the experiment was prepped in a BHI media with a total volume of 205 μL (200 μL of BHI, 5 μL of 1.5 OD cells) per well. Following 16 hours of growth at 37° C, OD at 600 nm (OD600) was measured with an Enspire Multimodal Plate Reader (Perkin Elmer). An IC_50_ measurement was taken for 12 colonies and each with 3 replicates.

In order to quantify the values for drug resistance, OD measurements for each drug concentration were normalized by OD measurements without the drug, and the dose-response curve was then fit to a Hill function (*g*(*d*) = (1 + (*d/k*)^*h*^)^−1^), with *d* the drug concentration, *k* the IC_50_, and *h* the Hill coefficient.

### Estimating growth costs

Growth costs were estimated by fitting time-series of optical density (for isolates grown in drug-free media) to a logistic growth function (***Zwietering et al., 1990***) given by *g*(*t*) = *g*_0_+*K* (1 + exp(4*μ*(*λ* − *t*)*/K* + 2))^−1^, where *μ* is the maximum specific growth rate, *λ* is the lag time, and *K* is the carrying capacity. The fitness cost (*c*) is then defined as *c* = 1 − *μ/μ*_*wt*_, where *μ*_*wt*_ is the maximum growth rate of the ancestral (WT) strain.

### Whole-genome sequencing

To see if there are any genomic changes that influence the measurement of collateral phenotypes, we sequenced 40 individual colonies that evolved DAP mutants and 2 V583 ancestors for a control. Illumina Short Read sequencing (400 Mbp / 2.7 million reads) and DNA isolation were performed by the Microbial Genome Sequencing Center (MiGS) at the University of Pittsburgh. We also performed population level sequencing (2000Mbp/13.3 Million reads) on samples drawn from days 2, 4, 6, and 8 from each population (performed at SeqCoast Genomics).

The resulting genomic data was analyzed using the high-throughput computational pipeline breseq (***Deatherage and Barrick, 2014***; ***Barrick et al., 2014***), with default settings (and using polymorphism mode for population sequencing). Briefly, genomes were trimmed and subsequently aligned to *E. faecalis* strain V583 (Accession numbers: AE016830 - AE016833) via Bowtie 2. A sequence read was discarded if less than 90 percent of the length of the read did not match the reference genome or a predicted candidate junction. At each position a Bayesian posterior probability is calculated and the log10 ratio of that probability versus the probability of another base (A, T, C, G, gap) is calculated. Sufficiently high consensus scores are marked as read alignment evidence (in our case a consensus score of 10). Any mutation that occurred in either of the 2 control V583 strains was filtered from the results.

For population sequencing, we removed any variants that did not occur at a frequency of greater than 25 percent on at least one day. For simplicity, in the figures in the main text we also removed variants if they were either 1) not observed in any single isolates and 2) had not previously been linked with DAP resistance. Detailed Muller plots containing these additional mutations is shown in Figure S1. To make Muller plots, we used the Lolipop pipeline, which is briefly described in (***Scribner et al., 2020***) and available on GitHub (https://github.com/cdeitrick/Lolipop).

### Long-read sequencing to confirm structural mutations

We confirmed the large-scale structural mutations in liaX arising in population DAP2 using longread Nanopore sequencing (***Deamer et al., 2016***). Genome assemblies and sequencing were performed by MiGS according to the following: Quality control and adapter trimming was performed with bcl-convert and porechop for Illumina and ONT sequencing respectively; hybrid assembly with Illumina and ONT reads was performed with Unicycler; assembly statistics were recorded with QUAST and assembly annotation was performed with Prokka (Default Parameters + ‘–rfam’).

### Near-perfect genome assembly combining long and short read sequencing

In general, we followed the method described in recent work on perfect bacterial genome assembly (***Wick et al., 2023***). We begun by using the Filtlong tool on our ONT long read sequencing data to remove reads shorter than 6000 base pairs, followed by discarding the worst 10% of long reads. We then used Trycycler to subsample 12 groups of semi-independent reads subsets for each population sequenced. Half of the 12 subsamples were then assembled used Raven, while the other half were assembled using Flye, resulting in 12 semi-independent genome assemblies. Tryclycler’s cluster function was used to cluster contigs accross the subsample assemblies and identify genuine replicons. This was followed with Trycycler’s reconcile command on each cluster and contigs unable to circularise were discarded. Trycycler’s msa (multiple sequence alignment) command was run on each cluster, followed by partitioning our filtered raw reads to a cluster. Finally, Trycycler consensus was run on each identified contig to generate a consensus sequence. The consensus sequence for each population was parsed via Medaka polishing to conservatively remove remaining errors in the long read assembly.

This process left us with an assembled long read sequence that is likely to be structurally accurate, but may contain small-scale errors such as single nucleotide errors or single bp deletions or insertions. We then leveraged Polypolish with the Illumina short reads to fix these small-scale errors, leaving us with a near-perfect genome assembly.

## Supplemental Information

This section contains supplemental figures referenced throughout the text

**Figure S1.**
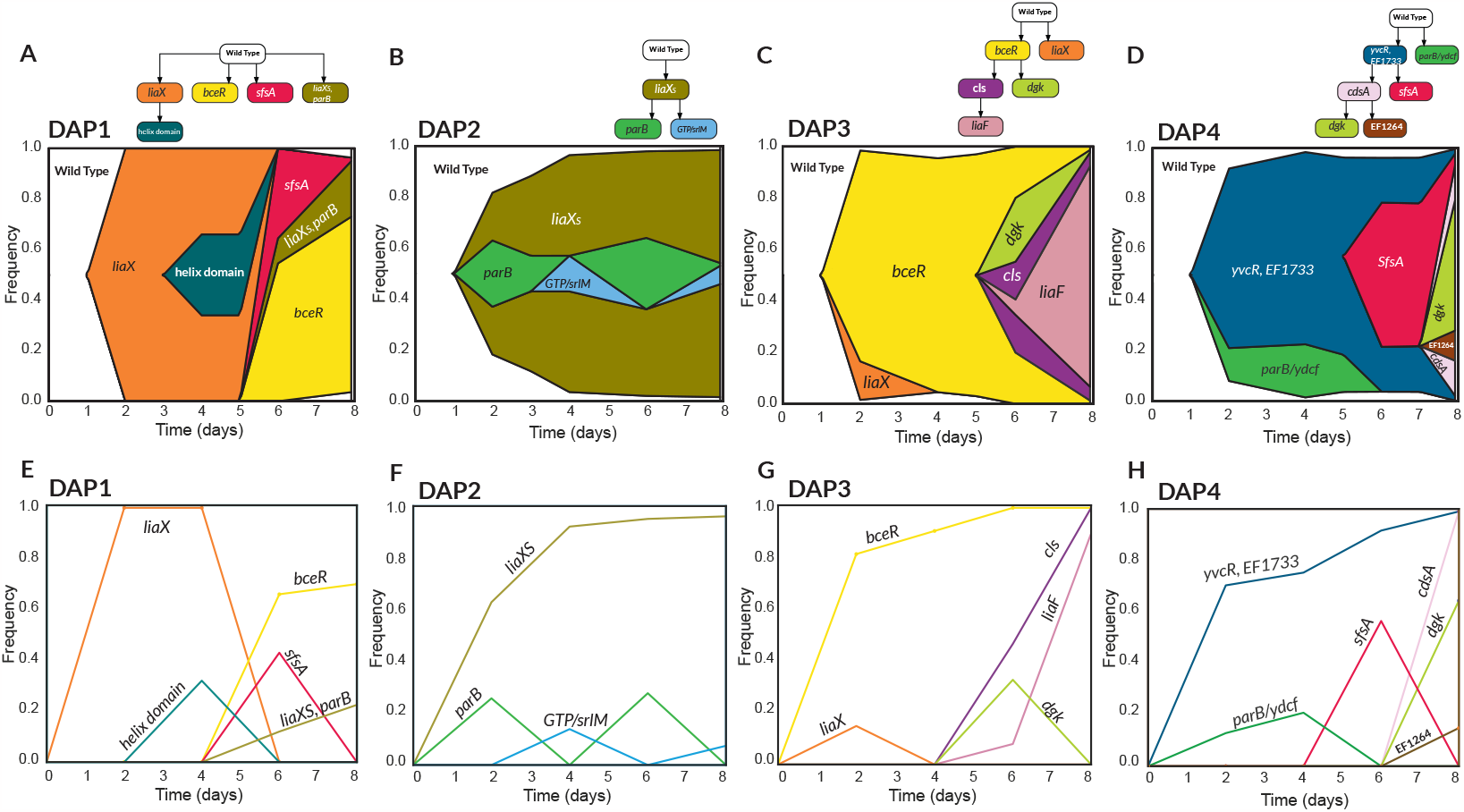
Muller plot from all mutations. Panels A–D correspond to the genotype tree and muller plots for each DAP population during evolution. Panel E-H plots individual genotype frequencies corresponding to each Muller plot.

**Figure S2.**
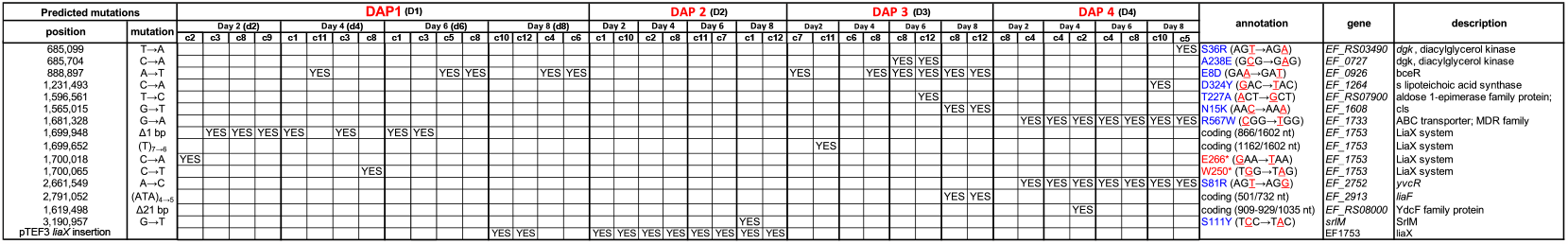
DAP mutaions. Mutations in 40 individual isolates from DAP1-DAP4 on different days.

**Figure S3.**
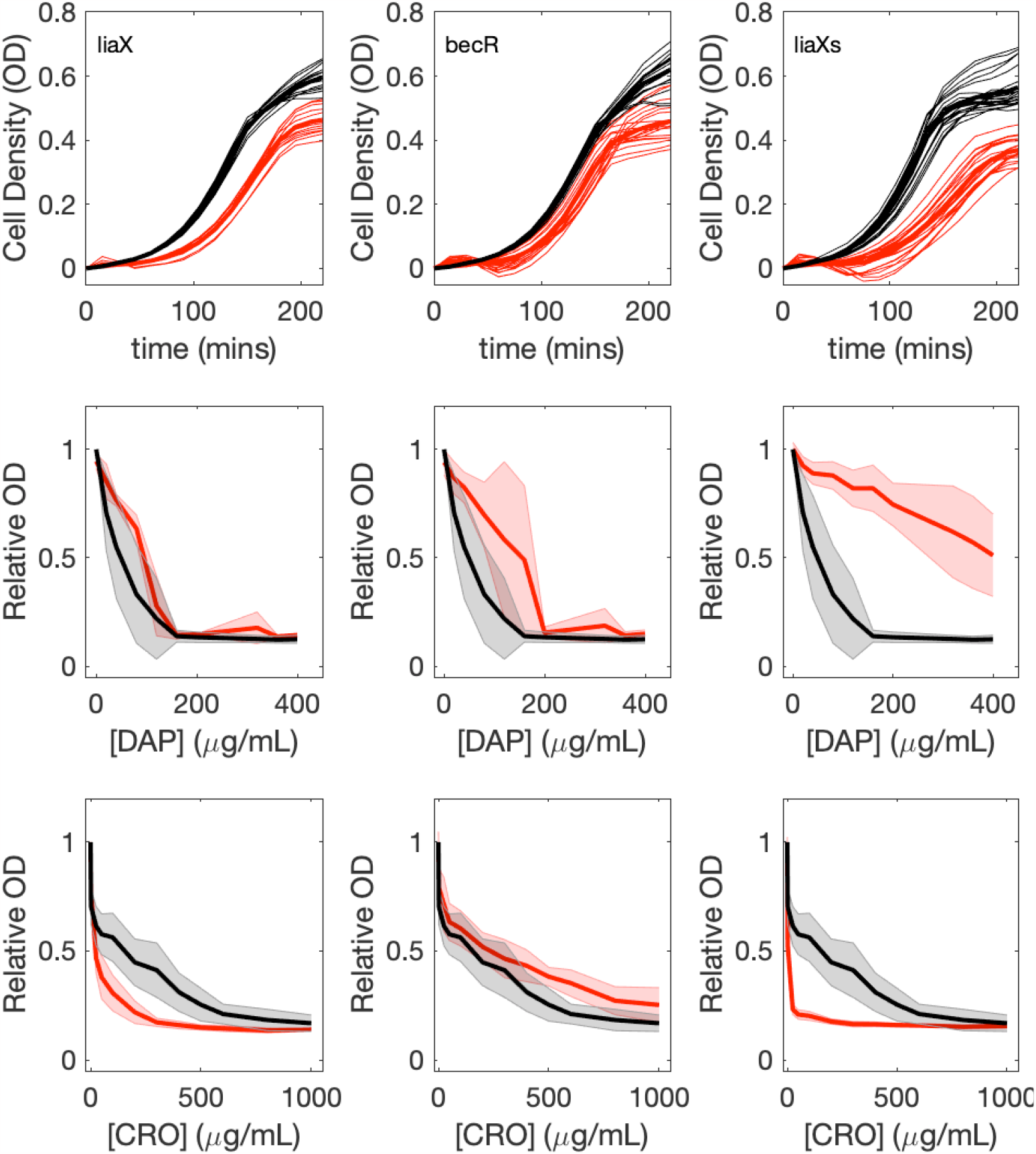
Drug-free growth curves and dose-response curves for single DAP-resistant mutants.

**Figure S4.**
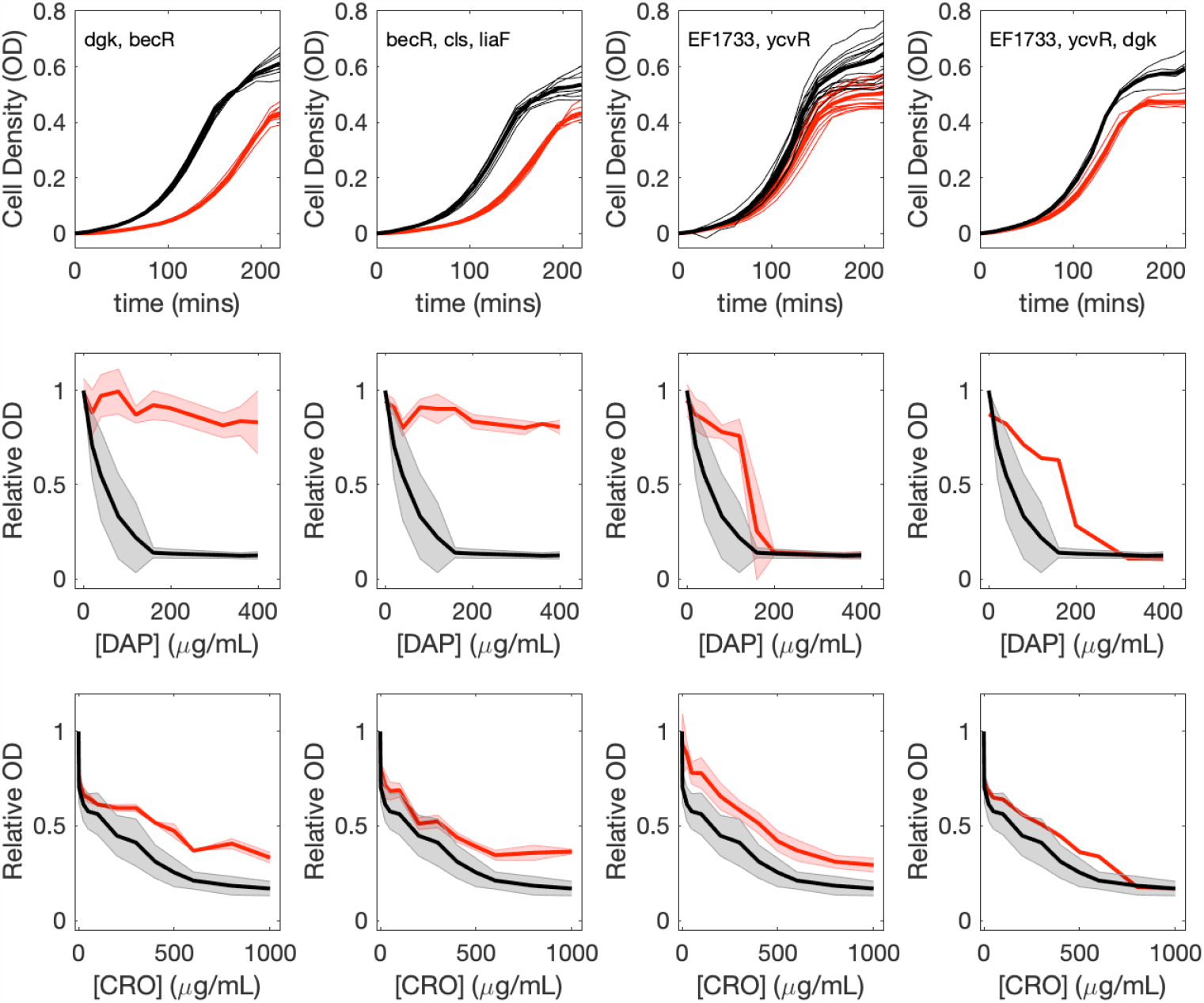
Drug-free growth curves and dose-response curves for double and triple DAP-resistant mutants.

**Figure S5.**
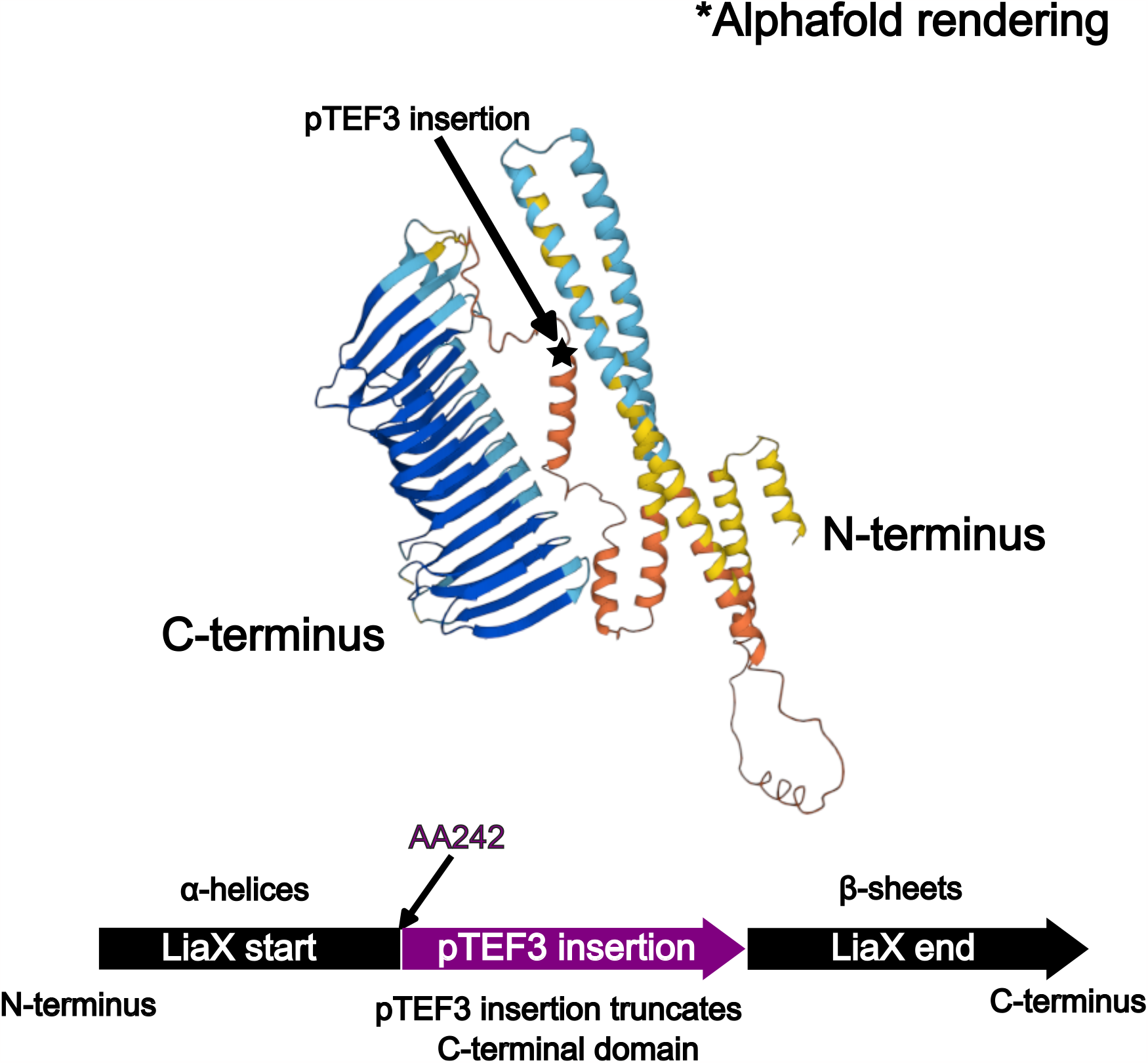
DAP mutaions. Alphafold rendering of the predicted molecular structure of *liaX*. Plasmid pTEF3 insertion occurs in a likely linkage region resulting in truncation of C-terminal domain.

